# Adhesion to a common ECM mediates interdependence in tissue morphogenesis in *Drosophila*

**DOI:** 10.1101/2025.03.15.643376

**Authors:** LE Sánchez-Cisneros, M Barrera-Velázquez, Dimitri Kromm, P Bun, H Merchant-Larios, LD Ríos-Barrera

## Abstract

For organs to be functional, the cells and tissues that constitute them must effectively interact with each other and coordinate their behaviours. Halfway during *Drosophila* embryogenesis, two lateral epidermal sheets stretch to fuse at the dorsal midline; concomitant with this, the main tubes of the respiratory system also shift dorsally. Here, we show that these processes occur simultaneously and are coordinated by the adhesion of the epidermal sheets and a subset of cells of the tracheal trunks to a common extracellular matrix (ECM) that separates them. We show that during dorsal closure, tracheal trunk cells extend protrusions towards the ECM underneath the epidermis. These protrusions are under tension, suggesting they have a mechanical function. Additionally, perturbing adhesion between tracheal cells and the epidermis affects the development of both tissues. Altogether, our findings uncover a novel mechanism used for tissue coordination during development, one that is based on tissue adhesion towards a common ECM capable of transmitting forces across the embryo.

## Introduction

Cells and tissues modify their behaviour according to extracellular signals. Many works have characterised the intracellular mediators and effectors of such signalling pathways and how they influence individual cell behaviour, leading to tissue-wide rearrangements. However, fewer studies have examined how seemingly independent tissues influence each other’s behaviours. The development of the *Drosophila* tracheal system is a very useful model for looking at tissue interactions and how they influence cell and tissue remodelling since tracheal tubes branch and migrate over neighbouring tissues of different biochemical and mechanical properties (Boube et al., 2001; Franch-Marro and Casanova, 2000; Ghabrial and Krasnow, 2006; Hayashi and Kondo, 2018; Urbano et al., 2011). The variety of environments that tracheal cells are exposed to, suggests that the tracheal system has wide plasticity to adapt to specific conditions.

Tracheal development is orchestrated by Branchless (Bnl), a member of the fibroblast growth factor (FGF) family. Bnl is expressed in various tissues within the embryo where it acts as a chemoattractant for tracheal tip cells. Initially, Bnl signals to tracheal pits on the lateral sides of the embryo, inducing primary branching and fusion to form the main tubes of the tracheal system, the dorsal trunks. Dorsal trunks are two multicellular tubes that early in development localize to the sides of the embryo. Bnl also controls branch outgrowth from these trunks: dorsal branches grow towards the dorsal midline, whereas ventral branches towards the nervous system and the gut [(Samakovlis et al., 1996; Sutherland et al., 1996), reviewed in (Hayashi and Kondo, 2018)]. It is in late tracheal development that trunks are displaced dorsally, however, the mechanisms behind trunk displacement remain unknown. Since trunk cells do not show hallmarks of Bnl signalling activation, and Bnl signalling becomes restricted to tracheal tip cells (Llimargas, 1999; Miao and Hayashi, 2016), it is likely that this repositioning is driven by trunk-extrinsic forces.

Trunk displacement coincides with branch outgrowth and epidermal dorsal closure, raising the question of whether forces from these processes contribute to trunk repositioning. During dorsal branch formation, tip cells migrate in response to Bnl, promoting stalk cell intercalation and branch elongation (Caussinus et al., 2008). Parallel to this the epidermis undergoes dorsal closure, a process in which the two lateral epidermal sheets stretch and fuse at the midline. The epidermis is pulled by the amnioserosa, an epithelium attached to the dorsal epidermis that generates force through actomyosin contraction (Duque and Gorfinkiel, 2016; Kiehart et al., 2000; Pasakarnis et al., 2016).

We show here that epidermal dorsal closure and dorsal branch migration participate in proper trunk displacement and morphogenesis. In addition, our experiments indicate that perturbing interactions between tracheal trunks and the epidermis also affects epidermal dorsal closure. Altogether, these results uncover a novel layer of regulation in tracheal morphogenesis, where multiple forces and adhesion to adjacent tissues mechanically couple the morphogenesis of multiple tissues.

## Materials and methods

### Fly lines

Fly stocks were kept under standard culture conditions. Experiments were carried out at 25°C or 29°C where stated. *btl-gal4, UAS-*mCD4::mIFP*; btl-gal4, UAS-*PH::mCherry*; btl-gal4, UAS-*Utrophin::GFP, and *Collagen-IV::GFP* were used before and come from Maria Leptin’s Lab at EMBL, Heidelberg (Ríos-Barrera and Leptin, 2022; Ríos-Barrera et al., 2017). *en-gal4, UAS*-GFP is from Tobias Troost, Heinrich Heine University Düsseldorf, Germany; *sqh-*Gap43::mCherry is from Thomas Lecuit lab, IBDM, France (Rauzi et al., 2010); *kay^1^*was obtained from Juan Riesgo-Escovar, UNAM, Mexico (Riesgo-Escovar and Hafen, 1997). Insertion of GFP at the *mys* locus (β-Integrin::GFP) was a kind gift from Nick Brown, MRC, UK (Klapholz et al., 2015) and insertion of the YFP derivative YPet at the *rhea* locus (Talin::YPet) is from Frank Schnorrer, IBDM, France (Spletter et al., 2018). *UAS-*CD8::GFP is from Fanis Missirlis, CINVESTAV, Mexico. The following lines come from Bloomington Drosophila Stock Center: *sr-gal4* (#26663); *UAS-mys* TRiP (#27735); *en-gal4, UAS-nls::mCherry* (#38420); *UAS-NsImb-vhhGFP4* (#38421); *UAS-Mmp1.f2* (#58702); *UAS-Mmp2* (#58705 and #58706); insertion of GFP at the *shg* locus (E-Cadherin::GFP, #60584); *UAS-Moe.myc.T559A* (*Moe^DN^;* #52234) *btl-moeRFP* (#64233); and *UAS-Dad* (#98452). To select genotypes of interest, we used balancers with fluorescent reporters from BDSC: TM6B,dfdYFP (#8704), CyO,sChFP and TM3,sChFP (both in #92597) *UAS-Talin* RNAi comes from the Vienna Drosophila Resource Center (#40399).

### Immunostainings

Immunostainings were carried out using standard methods. Embryos were dechorionated using 50% commercial bleach for 30 seconds, then washed with water and transferred to heptane. Afterward, embryos were fixed in 37% formaldehyde for 8 minutes. For devitellinization, formaldehyde was replaced with methanol, and vials were vigorously shaken for 1 minute. We rehydrated the embryos using PBS Triton X-100 0.3% and washed them three times. We then blocked using the washing solution and BSA 1%, and incubated overnight with the corresponding antibody at 4°C. The next day, we did three 10 min washes and incubated with secondary antibodies for 1 hour. These were then washed three times for 10 minutes each, and finally, we mounted them using Vectashield with DAPI. The antibodies used were: anti-GFP coupled to FITC (Abcam #ab6662; RRID:AB_305635, 1:500); anti-βIntegrin (Developmental Studies Hybridoma Bank [DSHB] #CF.6G11; RRID:AB_528310, 1:20); anti-Gasp (DSHB #2A12; RRID:AB_528492, 1:20) and as secondary antibodies, goat anti-mouse IgG coupled to Alexa647 (Thermofisher #A21235; RRID:AB_2535804, 1:300) and goat anti-mouse IgM coupled to Alexa647 (Thermofisher #A21238; RRID:AB_2535807, 1:300). Samples were analysed in a Nikon A1R+ confocal microscope using a resonant scanner and a 60x/1.2 NA water objective and processed using Fiji version 1.54p (Schindelin et al., 2012).

### Live microscopy

Embryos were dechorionated as above, washed with water and transferred to an agar plate to manually select the embryos of the desired stages. For confocal live imaging, embryos were glued to glass-bottom dishes using heptane glue and covered with halocarbon oil. Samples were analysed in a Nikon A1R+ confocal microscope using a resonant scanner and a 60x/1.2 NA water objective and processed using Fiji (Schindelin et al., 2012). Z-step size was 0.213 μm.

For light-sheet imaging, dechorionated embryos were mounted in low-melting temperature agarose within a glass capillary. The gel cylinder containing the embryo was then pushed above the glass rim to have optical access to it. The position of the embryo was then fixed, and the glass capillary was placed inside the PBS-filled imaging chamber of a homebuilt MuVi-SPIM. 3D volumes were acquired as described previously (Caroti et al., 2018; Medeiros et al., 2015), using two 20x/1.0 NA water immersion detection objectives (Olympus XLUMPLFLN20XW) for fluorescence detection and two 10x/0.3 NA water immersion objectives (Nikon CFI Plan Fluor 10X W) for excitation. Images were processed in Fiji (Schindelin et al., 2012). The time resolution was 2 minutes per frame and the z-step was 1.5 μm.

### Laser micro-dissection

Embryos were processed for live microscopy as above. Laser micro-dissection experiments were carried out as detailed previously (Sánchez-Cisneros et al., 2023). We used a Zeiss LSM 780 microscope, using a 63x/1.4NA oil objective. We used a femtosecond-pulsed two-photon laser at 950 nm, 75% (1540 mW) power. Micro-dissection was done using a single cycle on a single 1 μm thick z-stack.

### Larval fixations

Third-instar larvae were heat-fixed by transferring them to halocarbon oil in a cover slide and a heating plate at 65°C for 15 seconds. Afterward, the larvae were oriented with the dorsal side down. Larvae were analysed in a Nikon A1R+ confocal microscope using a resonant scanner in mosaic mode and a 10x/0.25 NA air objective and processed using Fiji.

### Electron microscopy

Embryos were processed according to the method of Tepass & Hartenstein (Tepass and Hartenstein, 1994). After dechorionation, the embryos were fixed in a glutaraldehyde 2% solution and heptane in a relation of 2:8 for 17 min in agitation. After fixed, we withdrew the vitelline membrane manually with insulin needles. Embryos were fixed in 2.5% glutaraldehyde in cacodylate solution (50mM, pH 7.2) and post-fixed in osmium tetroxide solution (2%). After fixations, the embryos were gradually dehydrated with alcohol solutions (70, 80, 90, 95, 100%) and acetone. The dehydrated embryos were incubated in acetone/Epon solution overnight and then were included in Epon solution for 2 days. Afterward, 1 μm semi-thin sections were stained with Methylene blue and visualized under a conventional light microscope to identify the regions of interest. Once these were identified, we did ultrafine (50-100 nm) sections and analysed them using transmission electron microscopy. This was carried out using a JEOL JEM 1200 EXII electron microscope. Feature tracing was done in Inkscape (version 1.3.2).

### Fluorescent *in situ* hybridization

We used the Hybridization Chain Reaction system (HCR RNA-FISH, Molecular Instruments) in an RNAse-free environment. Embryos were fixed in a mixture of heptane and 3.2% paraformaldehyde (PFA) in PBS for 30 minutes, and then devitellinized by removing PFA, adding methanol and shaking vigorously for 1 minute. Then, we washed them with ethanol and incubated overnight in 100% ethanol at 4°C. The next day we washed with 0.5% PBS Triton X-100 for 30 minutes and incubated in pre-warmed hybridization buffer at 37°C for 30 minutes. We incubated overnight in *kay* probe (1:250, designed to target all *kay* isoforms and for using with B3 amplifier) and probe solution at 37°C. Next, we washed with pre-warmed probe solution 4x, 15 minutes each, and washed twice with 5x SSCT 5 minutes at room temperature. Afterwards, we incubated in amplification buffer for 30 minutes at room temperature. We snap-cooled hairpins h1 and h2 separately (B3 amplifier labelled with Alexa594) by heating them at 95°C for 90 seconds and cooling to room temperature in the dark before mixing them with amplification buffer at room temperature. This was then added to embryos and incubated overnight at room temperature. Finally, the next day we washed with 5x SSCT at room temperature for 5, 5, 30 and 30 minutes and mounted using vectashield with DAPI. Images were acquired in a Nikon A1R+ confocal and a 60x/1.2 NA water objective.

### Image processing, analysis and statistics

We registered all time-lapse micrographs using the ‘Correct 3D drift’ function. Cartographical projections of the embryo surface were done using a custom-made Fiji macro that extracts the curvature of the embryo and reslices over 16 z planes throughout the time-lapse. To measure tissue coordination, we used particle image velocimetry (PIV) using the Fiji plugin described in (Tseng et al., 2012), with an interrogation window size of 128×128 px (6.25% of the image) with an overlap of 64 px, and a correlation threshold of 0.60. Cross-correlation analyses were done using a custom-made R code, using the average of magnitudes in the y-axis generated by the PIV plugin.

Embryonic and larval dorsal trunk length measurements were done manually in 3D using Imaris under a trial license, and these were normalised to the minimum distance between the initial and final measurement points. Embryonic dorsal trunk measurements were always done right after epidermal dorsal closure was completed. Rates of dorsal closure progression were measured using the Manual Tracking plugin from Fiji, following the signal corresponding to the leading edge for the epidermis, and a dorsal edge of the tracheal trunk. In both cases, we followed the same segment corresponding to the centre of the amnioserosa (Abdominal segments 3-4). We plotted displacement in the y axis, normalizing the data with the initial and final time points set to 0 and 100 (arbitrary units), respectively. To measure signal intensity in the deGradFP experiments, we did SUM projections and traced a line that crossed control and experimental regions (marked by the expression of *btl>*CD4::IFP or *en>*nls::mCherry) within each embryo. We plotted the fluorescence intensity within this line in both channels and calculated the correlation coefficient between the signal from the fluorescent reporter with respect to the signal of the target protein, β-integrin::GFP.

No blinding was carried out in any of our experiments. All our experiments represent pooled data from at least three independent crosses. We did not used statistical methods to determine sample size. All data were analysed and plotted using R Studio or Prism. For statistical analyses, data normal distribution was tested in Prism (version 10.5.0).

## Results

### The epidermis and tracheal trunks displace dorsally at the same time during embryogenesis

Halfway in development tracheal trunks displace from lateral to dorsal to acquire their final position, but the mechanisms behind this movement have not been described. Simultaneously with trunk displacement, dorsal branches elongate in response to Bnl (Figure 1A-A’’, Video S1). Only the tip cells respond to Bnl, and it was previously shown that force generated by tip cell migration allows stalk cell intercalation and branch elongation (Caussinus et al., 2008). However, whether the force generated by tip cell migration is sufficient to drive dorsal trunk displacement has not been determined. Nevertheless, at the same time as dorsal branches elongate, ganglionic and visceral branches extend ventrally and could also exert forces on the trunks (Figure 1B-B’’, Video S1). With branches growing out of the trunks in opposite directions, other factors likely participate in their dorsal displacement, but so far these remained unstudied.

**Figure 1.**
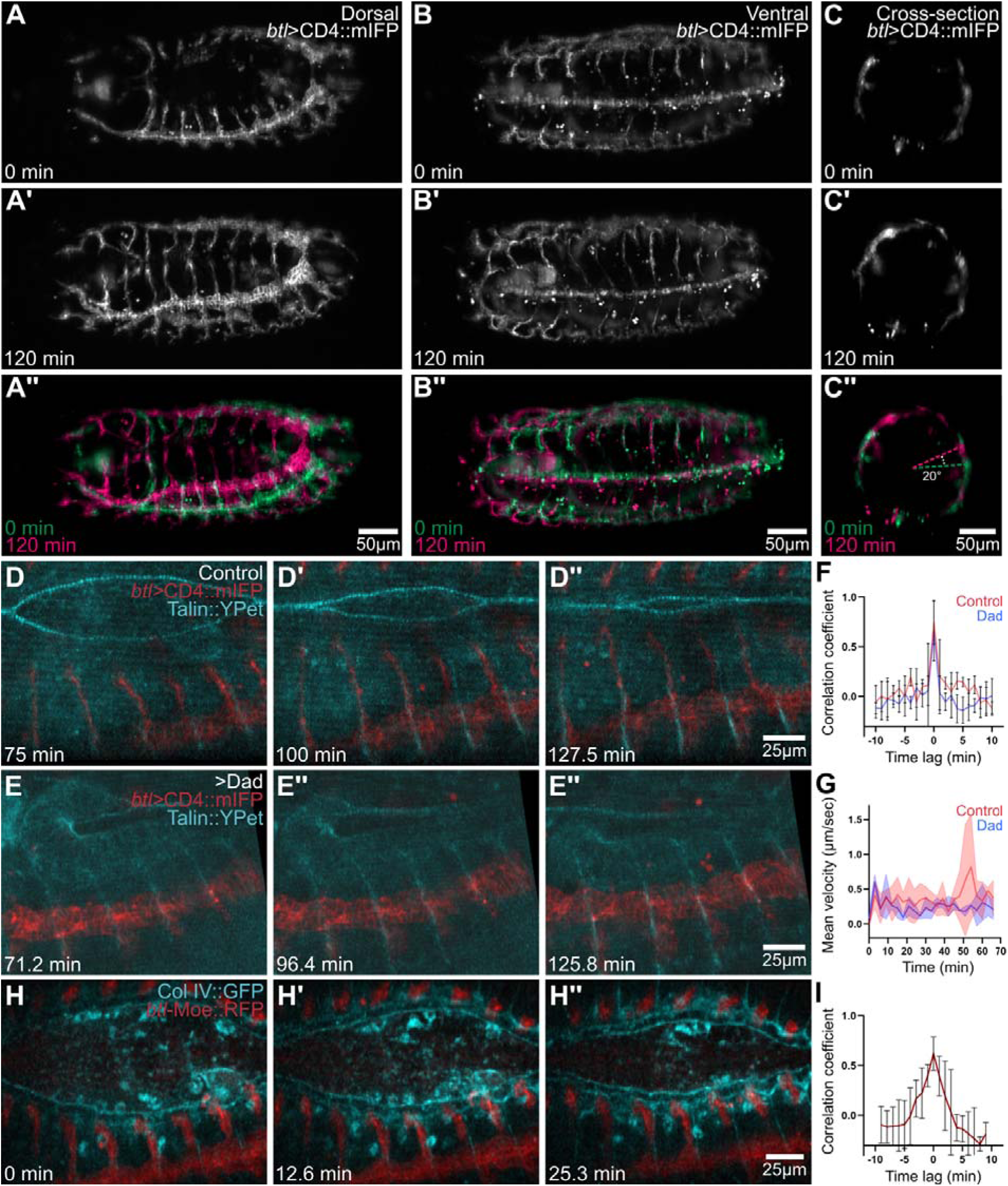
Coordination analyses of tracheal, epidermal and ECM movements during dorsal closure stages. (A – C’) Time-lapse imaging of tracheal development using MuVi-SPIM. The tracheal system is marked with *btl*>CD4::mIFP. (A – A’) Dorsal view; (B – B’) Ventral view; (C – C’) Transversal view. Cross-section was done between the 5^th^ and 6^th^ branch from anterior to posterior with respect to (A). (A’’ – C’’) Overposition of the initial (green) and final (pink) imaging time-point of the movie. (D – D’’) Cartographical maximum intensity projection of confocal stacks showing epidermis leading edge using Talin::YPet (cyan) and the tracheal system using *btl>*CD4::mIFP (red). (E – E’’) Embryo additionally overexpressing *UAS*-Dad. (F – F’’). Cross-correlation analysis of the signal displacement of Talin::YPet and *btl>*CD4::mIFP in Control (n=4) and Dad (n=5) embryos. (G) Velocity analysis of *btl>*CD4::mIFP signal in Control (n=3) and Dad (n=3) embryos. (H – H’’) Cartographical maximum intensity projection of confocal stacks showing Collagen IV::GFP and the tracheal system using *btl*-Moe::RFP. (I) Cross-correlation analysis of the signal displacement of Col IV::GFP (cyan) and *btl*-Moe::RFP (red) throughout time, n=5. Plots show means +/- SD.

Tracheal trunk displacement occurs at similar stages as epidermal dorsal closure, so we wondered if these two processes could be actively coordinated, and if the epidermis could participate in trunk displacement. To study this, we first carried out double-tissue labelling and live imaging following the Moesin actin-binding domain fused to a red fluorescent protein expressed under direct control of the tracheal-specific promoter of *btl* (*btl-*Moe::RFP). We used the Gal4*/UAS* system to label spaced epidermal stripes using GFP (*en>*GFP). In these recordings, we confirmed that trunk displacement coincided with epidermal dorsal closure (Figure S1A-A’’). We quantified this observation by doing PIV analyses over consecutive timepoints for each of the tissues, followed by cross-correlation analysis of both tissue displacements. We found a very strong positive correlation (Figure S1B), suggesting that epidermal dorsal closure and trunk displacement are actively coordinated. We corroborated these measurements by following Talin endogenously tagged with YPet at the C-term (Talin::YPet) and an infrared fluorescent protein fused to CD4 under *btl-gal4* (*btl>*mCD4::mIFP), finding similar results (Figure 1D-D’’, F, Video S2).

Since dorsal branches elongate at the same time as dorsal closure, and tip cells migrate right behind the leading edge of the epidermis, it was still possible that trunk displacement was coordinated with epidermal dorsal closure indirectly, as consequence of branches pulling on the trunks as they migrate behind the epidermis leading edge. To test the role of dorsal branch migration in trunk displacement, we over-expressed the inhibitory SMAD Daughters against dpp (Dad) in the tracheal system using *btl-gal4*. Dad over-expression prevents Decapentaplegic (Dpp) signalling activation, a condition where dorsal branches are not formed (Ribeiro et al., 2004). We found that in the absence of dorsal branches, the trunks still moved dorsally in coordination with the epidermis and at the same velocity as controls (Figure 1E-G, Video S2). This supports that dorsal branches are dispensable for this movement and in addition, it suggests that ventral branches do not pull on the trunks, as in the absence of dorsal branches we still see the trunks moving dorsally.

The dorsal displacements of the trunks and the epidermis could be controlled by extracellular signals that act on both tissues. However, while signals from the epidermis participate in tracheal development, these act mainly in fate determination or tip cell migration (Affolter et al., 1994; Chihara and Hayashi, 2000; Glazer and Shilo, 2001; Kato et al., 2004; Llimargas, 2000). Similarly, the signals that regulate epidermal dorsal closure do not participate in tracheal development; namely, JNK signalling (Letizia et al., 2023). Therefore, we hypothesized that coordinated displacement could be mediated by adhesion, as the tracheal system lies beneath the epidermis. Tissue adhesion could be mediated by an intermediate extracellular matrix (ECM) layer. To explore this, we did time-lapse microscopy of the tracheal system together with an ECM marker, an endogenous insertion of GFP to the *viking* locus, which codes for Collagen IV α2 subunit (from now on referred to as Col IV::GFP, Figure 1H-H’’). Col IV::GFP was enriched surrounding cardioblasts, haemocytes, and at muscle attachment sites as reported previously (Borchiellini et al., 1996). Like what we found with epidermal markers, the signal displacements of Col IV::GFP and the tracheal reporter were positively correlated (Figure 1I).

### The tracheal trunk cells contact tendon cells of the epidermis during dorsal closure

Since we could not detect a particular enrichment of Col IV::GFP around the trunks using live imaging, we stained embryos against β-integrin and against GFP to detect Col IV::GFP. We could not detect any enrichment for these components at the tracheal trunks during the stages of dorsal closure but only after this process is completed, as had been reported already [Figure S1C-E; (Klußmann-Fricke et al., 2022)]. Instead, at the onset of dorsal closure, Col IV::GFP and β-integrin were only detectable at muscle attachment sites (Figure S1D-D’’).

Muscles attach to epidermal tendon cells through a specialized meshwork of ECM (Fogerty et al., 1994; Prokop et al., 1998; Urbano et al., 2011); we figured that tracheal cells could also interact with the epidermis at these points. Classical anatomical descriptions of the embryo report that muscles separate the tracheal trunks from the epidermis (Hartenstein, 1993). Nevertheless, we decided to obtain higher resolution images of these regions by using transmission electron microscopy. With this approach, we observed that as reported, the dorsal trunks and the epidermis are separated by muscles and other mesenchymal cells in most parts of the embryo (Figure 2A, S2A). However, in the regions where tendon cells are found, trunks are no longer separated from the epidermis by other tissues but are instead found adjacent to the epidermis (Figure 2B-B’’, S2B-B’’). Tendon cells are easily recognizable in these images as they localise at the segmental borders and associate to muscles. We identified tracheal cells based on their epithelial morphology and the presence of a lumen within the epithelium (Figure. 2C-C’’, S2C-E’). These experiments revealed a novel anatomical contact between the epidermis and tracheal trunks that had not been previously reported and whose relevance had not been studied up to now.

**Figure S1.**
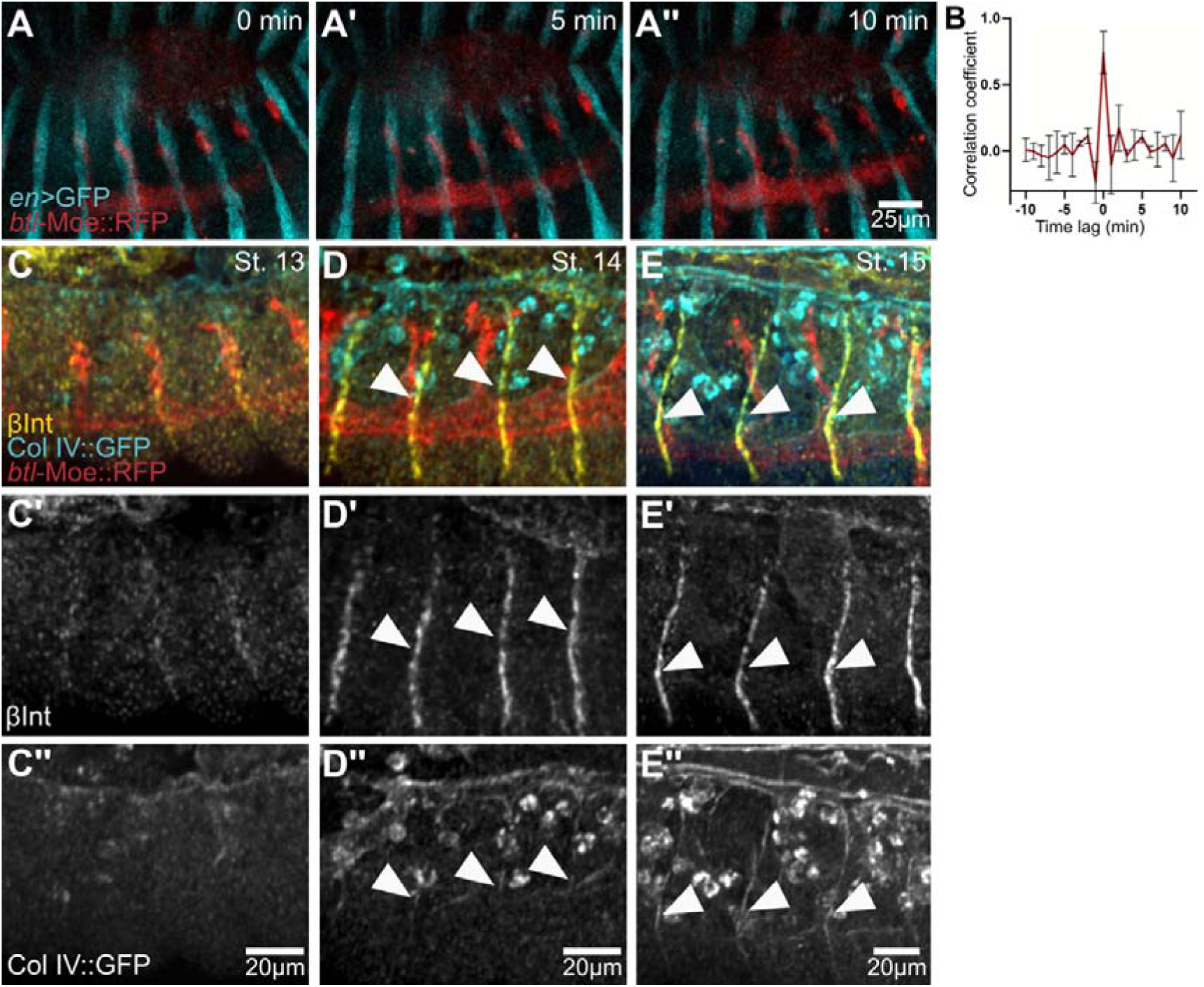
Analysis of epidermis-trachea coordination and distribution of Collagen IV::GFP and β-Integrin. (A – A’’) Cartographical maximum intensity projection of confocal stacks showing stripes of epidermis using *en>*GFP (cyan) and the tracheal system using *btl*-Moe::RFP (red). (B) Cross-correlation analysis of *en>*GFP and *btl-*Moe::RFP signal displacement. (C – E) Distribution of Collagen IV:: GFP (cyan) and β-Integrin (yellow) in stages 13-15 in relation to the tracheal system labelled with *btl-*MoeRFP (red). (A-A’’) Stage 13; (B-B’’) Stage 14; (C-C’’) Stage 15. Arrowheads point to muscle attachment sites at their intersection with tracheal trunks.

**Figure 2.**
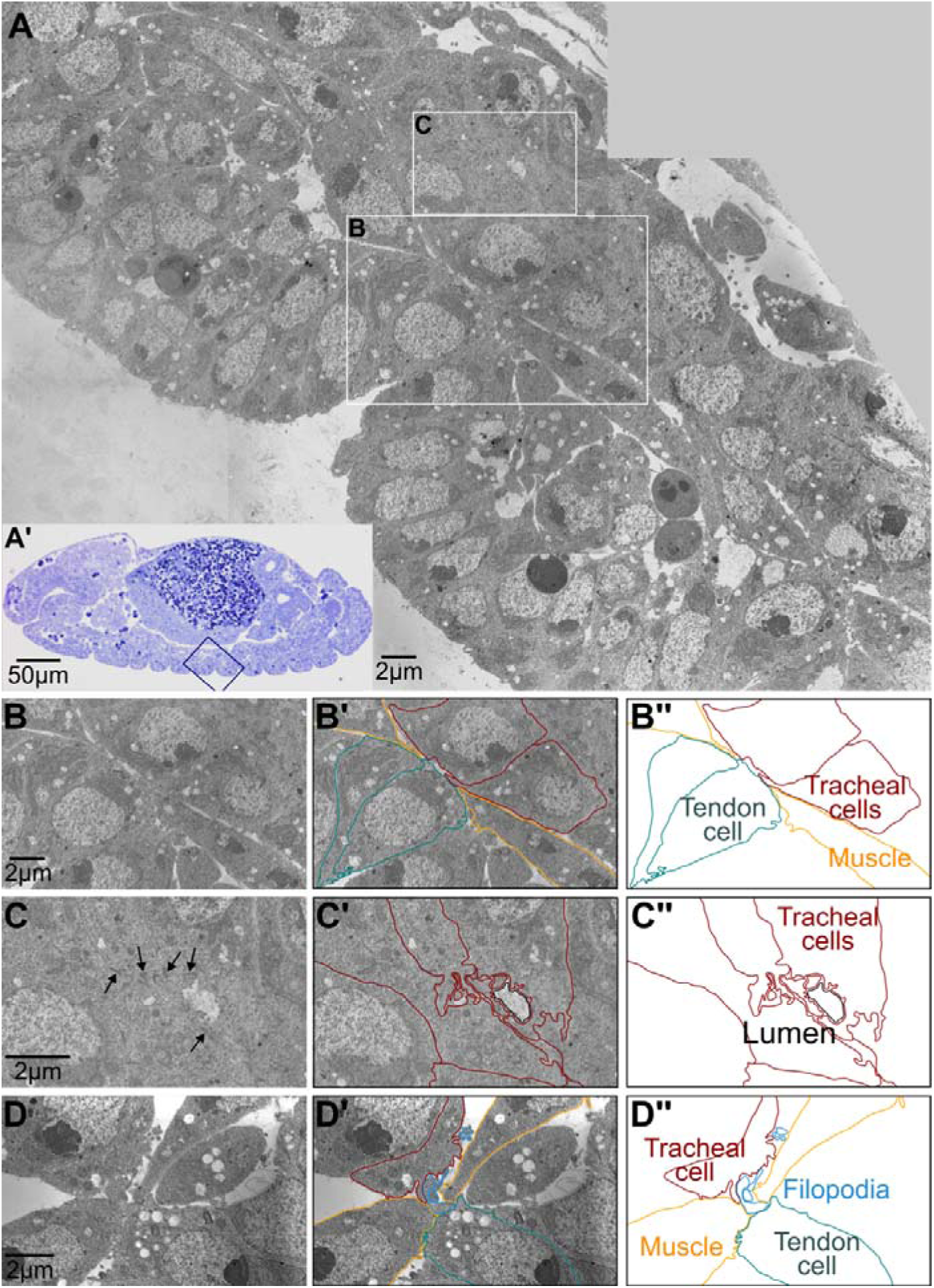
Contact points between the epidermis and the tracheal trunks. (A) Electron micrograph of a longitudinal section of the dorsal trunk, stage 14 embryo. The regions indicated by the white squares are magnified in (B-C). (A’) Semi-thin histological section used to identify the region of interest presented in (A), see methods. The squared area corresponds to the electron micrograph in (A). (B) Magnification of a contact point between tracheal trunks and a tendon cell. (B’, B’’) Manual tracings of the elements present in (B). (C) Magnification of a region of the tracheal trunk where the lumen is visible, arrows point to electron-dense areas that correspond to adherens junctions. (C’, C’’) Manual tracings of the elements present in (C). (D) Electron micrograph of tracheal cells in proximity to tendon cells and muscles. (D’, D’’) Manual tracings of the elements present in (D).

**Figure S2.**
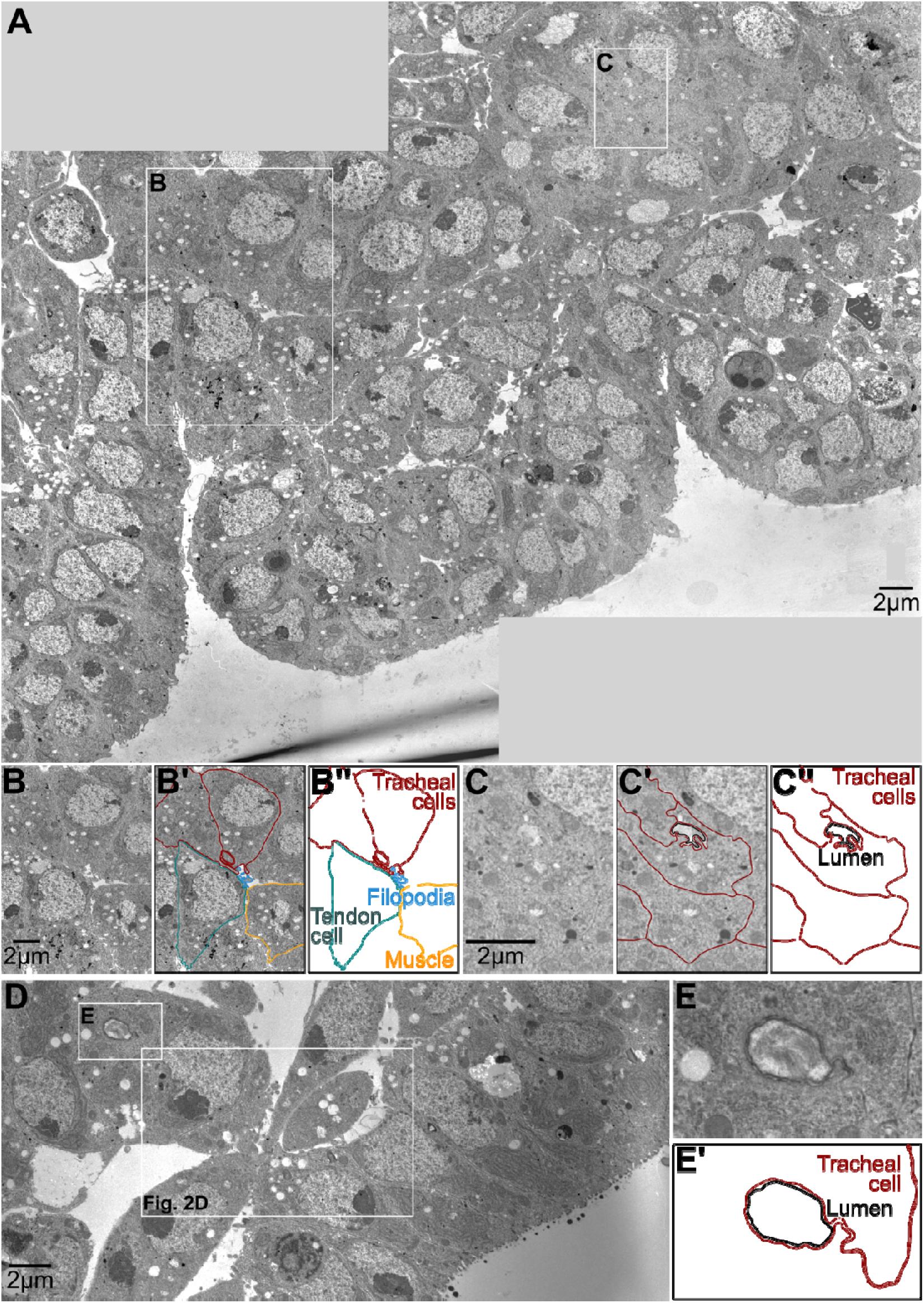
Contact points between the epidermis and the tracheal trunks (other examples). (A) Electron micrograph of a longitudinal section of the dorsal trunk, stage 14 embryo. The regions indicated by the white squares are magnified in (B-C). (B) Magnification of a contact point between tracheal trunks and a tendon cell. (B’, B’’) Manual tracings of the elements present in (B). (C) Magnification of a region of the tracheal trunk where the lumen is visible. (C’, C’’) Manual tracings of the elements present in (C). (D) Overview image of the micrograph magnified in Figure 2D. (E) Magnified view of the tracheal lumen illustrated in (D). (E’) Manual tracings of the elements present in (E).

### A subpopulation of dorsal trunk cells forms tensile protrusions towards the epidermis

Electron micrographs also revealed high filopodial abundance at the interphase between tendon cells and tracheal trunks (Figure 2D, S2B). Muscles extend filopodia as they attach to tendon cells (Maartens and Brown, 2015), leading us to examine whether tracheal cells could also form filopodia to make contact with tendon cells. Time lapse imaging using MuVI-SPIM suggested protrusive activity at tracheal trunks during dorsal closure stages (Figure 3A, Video S3). We performed high resolution live imaging of dorsal trunk cells to better visualize this protrusive behaviour. For this, we used *btl*>CD4::mIFP to label the tracheal system and a ubiquitously expressed membrane marker, *sqh*-Gap43::mCherry to label the epidermis. In time-lapse movies, we observed that trunk cells form protrusions in the direction of the epidermis (Figure 3B, Video S4). These protrusions are very stable; they persist in high time resolution acquisitions and throughout epidermal dorsal closure (Figure 3C, Video S5). Notably, the cells that form protrusions are evenly distributed at the trunks, near the base of each dorsal branch and at the intersection with muscle attachment sites (Figure 3A, B, Video S3). Also, they are still present in conditions of Dad over-expression, where dorsal branches are not formed (Figure 3D, Video S6). We propose that this subset of tracheal cells mediates the interaction between the trunks and tendon cells, and because of their morphology, we refer to them as protruding cells.

**Figure 3.**
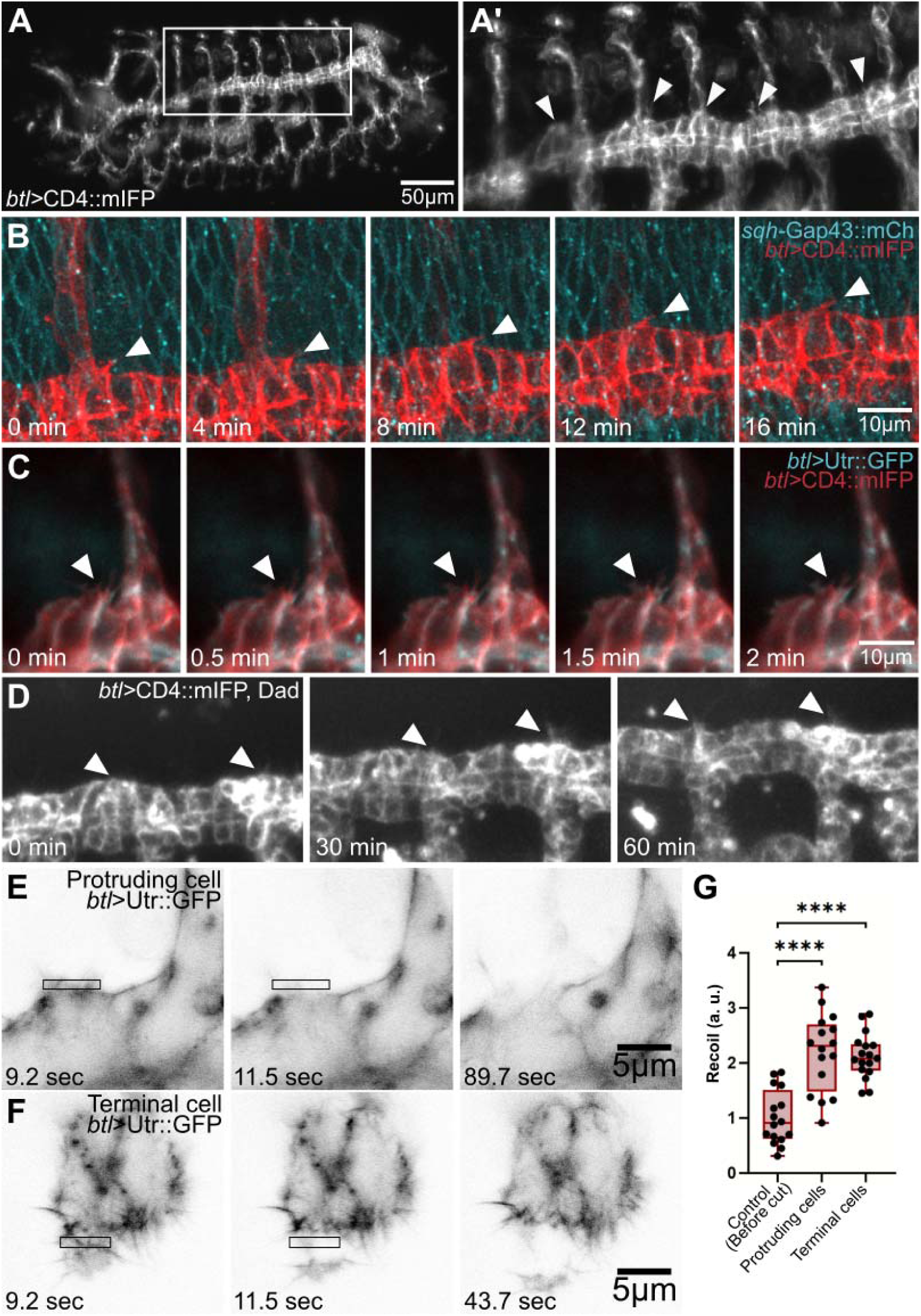
Live imaging of protruding cells and laser micro-dissection experiments. (A-A’) Time-lapse imaging of an embryo expressing CD4::mIFP under *btl-gal4* and imaged using MuVi-SPIM. (B) Time-lapse imaging of an embryo expressing CD4::mIFP under *btl-gal4* (red) and Gap43::mCherry under direct control the *sqh* promoter (cyan) and imaged using confocal microscopy. (C) Time-lapse imaging of an embryo expressing CD4::mIFP (red) and Utr::GFP (cyan) under *btl-gal4*. (D) Time-lapse imaging of an embryo expressing CD4::mIFP and Dad. Arrowheads in (A-D) point to protruding cells of the dorsal trunks. (E) Laser cut in the protruding cells of the dorsal trunk expressing *btl*>Utr::GFP. (F) Laser cuts in the terminal cells of the dorsal branch expressing *btl*>Utr::GFP. Laser cuts were done in the regions marked with black squares in (E – F). (G) Quantification of the recoil using PIV analysis. Control vs Protruding cells, p<0.0001; Control vs Terminal cells, p<0.0001. Control, n=16; Protruding cells, n=16; Terminal cells, n=17. Box plot represents median, interquartile range (IQR) and min and max values. Significance was determined using ANOVA and Dunnett correction for multiple comparisons.

If protruding cells adhere to the ECM of muscle-attachment sites, we hypothesised that they should be under tension. We tested this by doing laser micro-dissection directly over these structures and measuring the resulting recoil speed using PIV analyses. We found that seconds after doing the laser cut there was a significant recoil in the direction of the trunk (Figure 3E, G; Video S7), which was comparable to the recoil seen upon laser micro-dissection of tracheal tip cells [Figure 3F, G, Video S7; (Ríos-Barrera and Leptin, 2022)]. These results show that protruding cells are under tension, likely due to their role as mediators of trachea-epidermis interactions.

### Integrin-ECM adhesion complexes participate in the interaction between dorsal trunks and the epidermis

To test the relevance of the adhesion between protruding cells and muscle-attachment sites, we combined *btl-*Moe::RFP to follow the tracheal system, together with a tendon cell driver (*stripe; sr-gal4*) that allowed us to visualise these cells and carry out ECM perturbation experiments. We over-expressed matrix metalloproteinases (Mmp1 and Mmp2), enzymes that degrade ECM components (Jia et al., 2014; Llano et al., 2002), and observed their effect on tracheal development. In control embryos we observed evenly distributed stripes of tendon cells across the anterior-posterior axis of the embryo that traversed perpendicularly the tracheal trunks (Figure 4A – A’). In embryos where we over-expressed Mmp1 or Mmp2 in tendon cells, the general organization of tendon cells remained the same and tracheal trunks were still displaced dorsally; however, they developed tortuosities over time, a morphology that is not seen in control embryos of similar developmental stages (Figure 4B-C). Consequently, experimental embryos showed significantly longer dorsal trunks compared to controls (Figure 4D). As Mmp2 is a membrane-anchored protein (Llano et al., 2002) its protease activity is limited to the ECM adjacent to the tendon cells, reducing the possibility that Mmp2 expressed by the tendon cells could have a direct effect on tracheal trunks by diffusing through the extracellular space.

**Figure 4.**
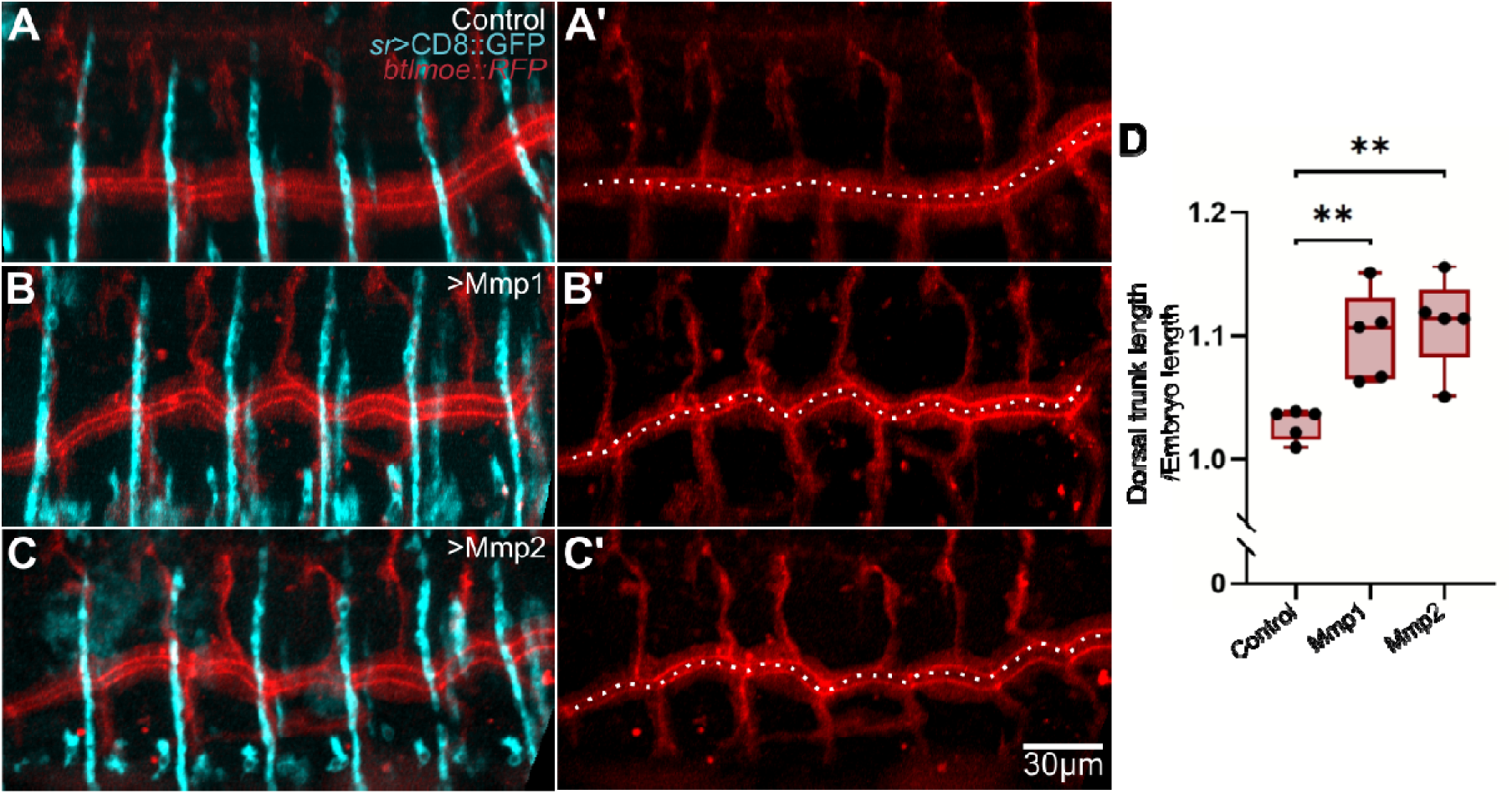
Effect of over-expressing MMPs in tendon cells on tracheal trunk morphology. (A-C) Live-imaging of embryos expressing CD8::GFP under the tendon cell driver *sr-gal4* (cyan) and *btl-*Moe::RFP (tracheal system, red). (A) Control; (B) Embryo additionally over-expressing *UAS-*Mmp1; (C) Embryo additionally over-expressing *UAS-*Mmp2. (D) Normalised dorsal trunk (DT) length. Control vs Mmp1, p=0.0067; Control vs Mmp2, p=0.0025. Box plot represents median, IQR and min and max values. Control, n=5; Mmp1, n=5; Mmp2, n=5. Significance was determined using ANOVA and Dunnett correction for multiple comparisons.

We wondered if the tracheal defects caused by Mmp2 expression in tendon cells would persist in later stages of development, so we looked at tracheal trunks in third instar larvae. We found a similar phenotype to the one we observed during embryogenesis, with dorsal trunks acquiring a wavy phenotype resulting in significantly longer trunks compared to controls (Figure S3A, C). These results show that perturbing the ECM that lies below the tendon cells influences tracheal trunk morphology.

**Figure S3.**
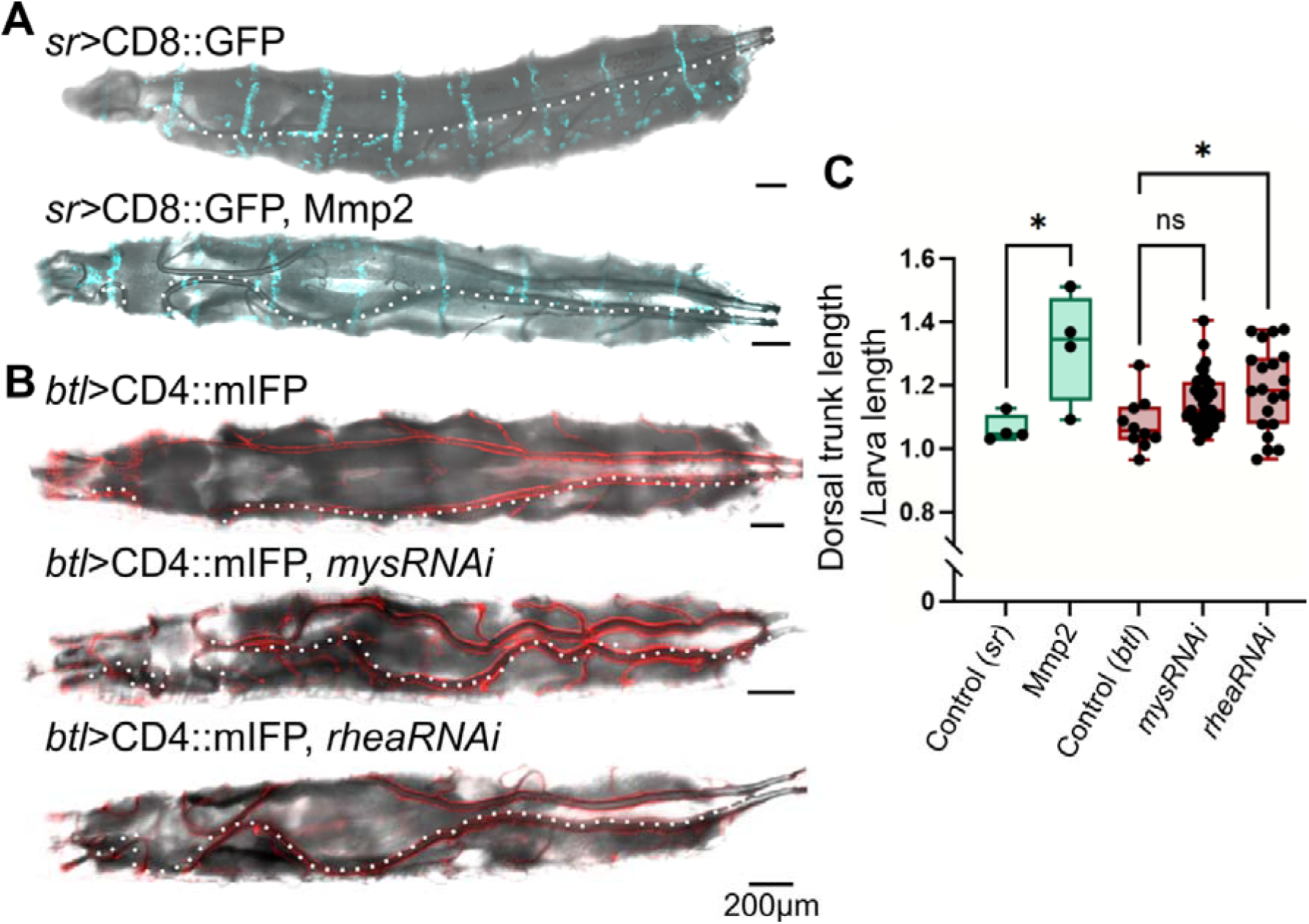
The role of adhesion complexes and ECM in larval dorsal trunk morphology. (A-B) Overlay of maximum-intensity projections of fluorescent reporters and minimum-intensity projections of reflected light images (for bright field) of heat-fixed third instar larvae. (A) Larvae expressing GFP under *sr-gal4*. Top panel, control; bottom panel, over-expression of Mmp2. (B) Larvae expressing CD4::mIFP under *btl-gal4*. Top panel, control; middle panel, *mys* (which codes for β-integrin) RNAi; bottom panel, *rhea* (which codes for Talin) RNAi. (C) Normalised dorsal trunk (DT) length. Control vs Mmp2, p=0.0204; Control vs *mysRNAi*, p=0.1057; Control vs *rheaRNAi*, p=0.0283. Box plot represents median, IQR and min and max values. Control (*sr*), n=4; Mmp2, n=4; Control (*btl*), n=10; *mysRNAi*, n=33; *rheaRNAi*, n=20. Significance was determined using ANOVA and Kruskal-Wallis test for multiple comparisons.

As a complementary approach, we also removed β-Integrin specifically in the tracheal system. Studies on integrins in the embryonic tracheal system have been hampered by abundant maternal contribution that may obscure early phenotypes, and/or by the early lethality of germ-line clones for mutations on components of the integrin complex (Brown, 1994; Wieschaus and Noell, 1986). Therefore, we used the deGradFP system that targets GFP-labelled proteins to proteasomal degradation. This is achieved by expressing UAS-NsImb::Vhh4, a modified ubiquitin ligase (NsImb) fused to a nanobody against GFP (Vhh4) that can be expressed under any particular driver (Caussinus et al., 2011). For these experiments, we targeted an endogenous insertion of GFP in the gene *mys*, that codes for β-Integrin (Klapholz et al., 2015). β-Integrin::GFP also allowed us to visualise the epidermis and other tissues in time-lapse experiments. Since β-Integrin::GFP is a transmembrane protein, it was possible that it could be unsensitive to proteasomal degradation using deGradFP. We tested the capacity of the system to degrade β-Integrin::GFP with *en-gal4* in stage 12-13 embryos and found that the β-Integrin::GFP signal significantly decreased in the regions were *en-gal4* was expressed (Figure S4A-B, E). Testing this using *btl-gal4* was technically more challenging, since by the time *btl-gal4* is active, the β-Integrin::GFP signal is more abundant in other tissues. Nevertheless, we also found a significant decrease in the GFP intensity (Figure S4CD, F).

**Figure S4.**
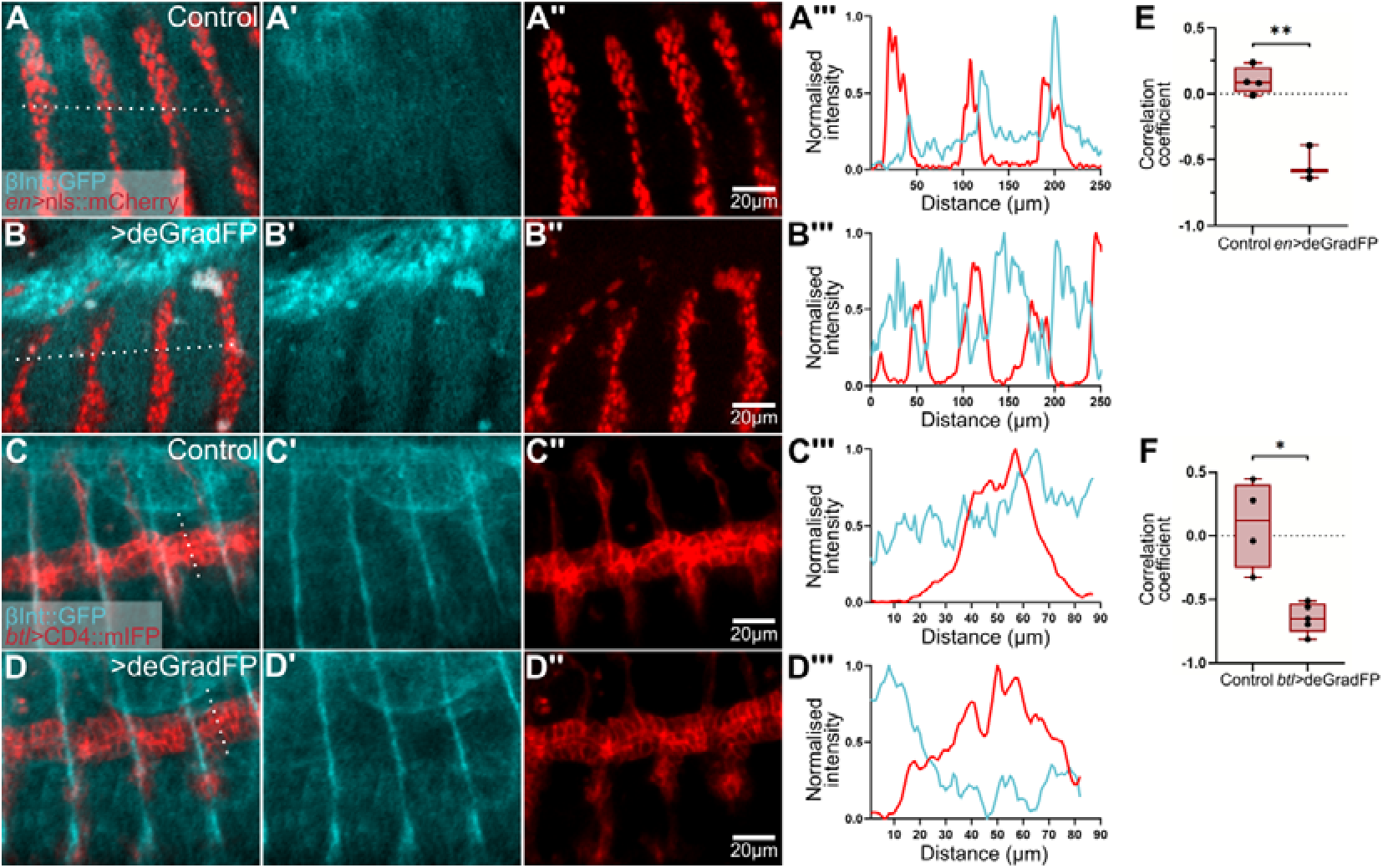
Effect of deGradFP expression on β-Integrin::GFP fluorescence intensity. (A-D) Maximum intensity projections of embryos expressing β-Integrin::GFP (cyan). (A-B) Embryos additionally expressing nls::mCherry (red) under *en-gal4*. (C-D) Embryos additionally expressing CD4::mIFP under *btl-gal4*. (A, C) Controls; (B, D) deGradFP over-expression. (A’’’-D’’’) Fluorescence intensity profiles across the dotted white lines drawn in (A-D). (E-F) Correlation analyses of fluorescence intensity profiles comparing signals from the *en>*deGradFP (E; p=0.0028; Control, n=4; *en*>deGradFP, n=3) and the *btl>*deGradFP (F; p=0.0185; Control, n=4; *btl*>deGradFP, n=5) experiments. Box plots represent median, IQR and min and max values. Significance was determined using t-tests.

When doing time-lapse microscopy, in control embryos we observed that tracheal trunks move to the dorsal side of the embryo in coordination with the epidermis, as determined by cross-correlation analyses (Figure 5A-C, G, Video S8, left). When we degraded β-Integrin::GFP in the tracheal system, as in our Mmp experiments, we observed coordinated tissue displacement (Figure 5D-G, Video S8, right) and tracheal trunks also showed a wavy lumen phenotype, resulting in significantly longer trunks (Figure 5H). We corroborated these results in third instar larvae using RNAi against *mys* and *rhea* (which code for β-Integrin and its intracellular mediator Talin, respectively). As with Mmp2 over-expression using *sr-gal4*, silencing these genes under *btl-gal4* resulted in wavy dorsal trunks (Figure S3B-C). These manipulations led to partial lethality (32% of expected animals hatched as adults, n=61), suggesting partial silencing or backup mechanisms that supress lethality. Nevertheless, these results show that cell-ECM interactions are important for proper trunk morphology during embryonic and larval development.

**Figure 5.**
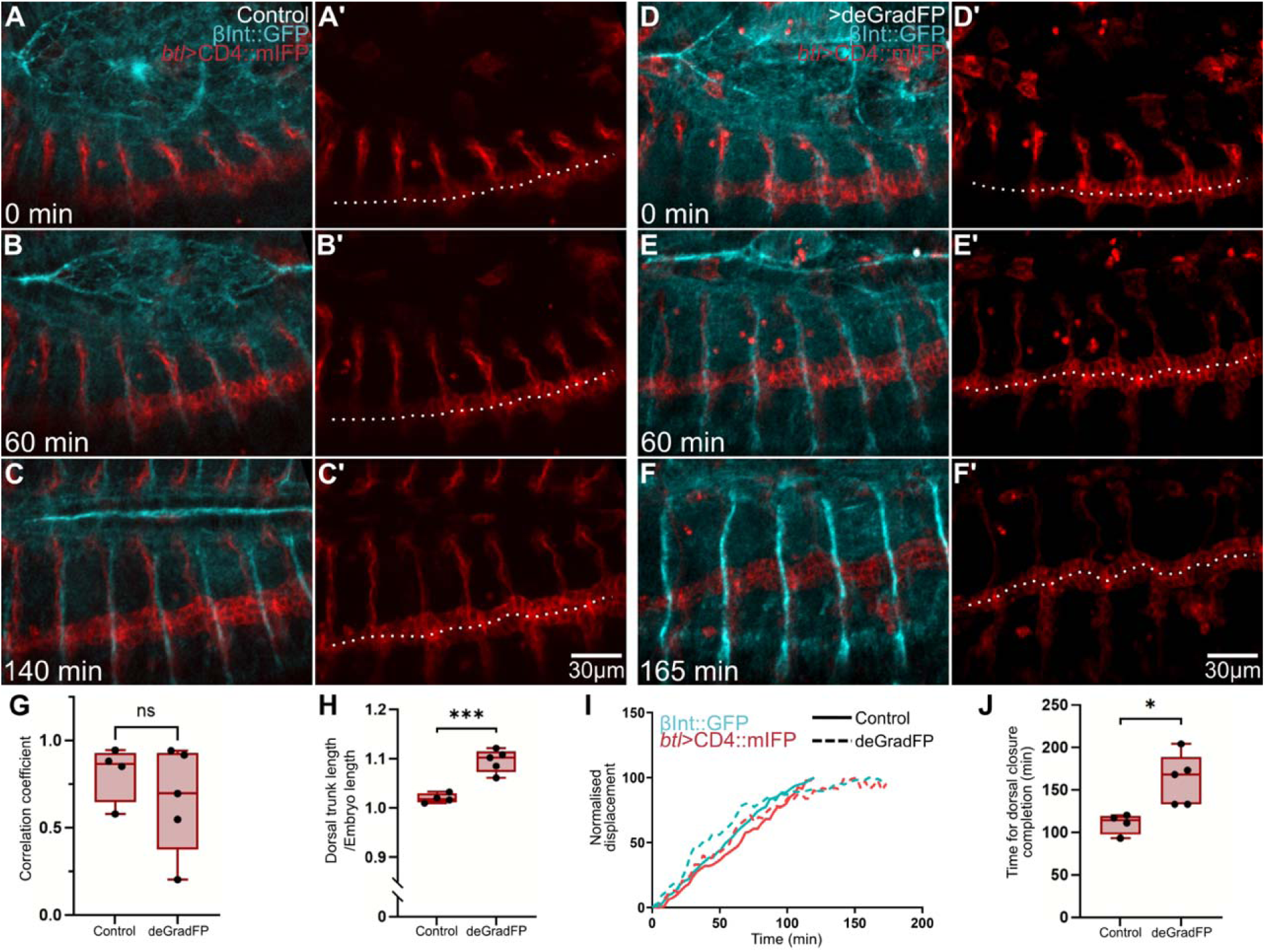
Effect of deGradFP expression in tracheal morphology of β-Integrin::GFP embryos. (A-F) Maximum intensity projections and time-lapse imaging of embryos expressing β-integrin::GFP (cyan) and CD4::mIFP under *btl-gal4* (red). (A-C) Control; (D-F) Embryo additionally expressing *UAS-*deGradFP. Dotted lines in (A’-F’) highlight the tracheal trunk lumen. (G) Quantification of epidermis-dorsal trunk coordination using PIV and cross-correlation analyses, p=0.3674. (H) Normalised dorsal trunk (DT) length, p=0.0008. (I) Normalised displacement of epidermis (cyan) and dorsal trunk (red). (J) Time for dorsal closure completion, p=0.0144. Control, n=4; deGradFP, n=5. Significance was assessed using t-tests. Box plots in (G, H, J) represents median, IQR and min and max values.

### Affecting dorsal closure results in aberrant tracheal morphogenesis and vice versa

In our deGradFP experiments, we noticed that in addition to the effects on tracheal trunks, epidermal dorsal closure also appeared to be affected upon depletion of β-Integrin::GFP from the tracheal system (Figure 5F, Video S7, right). We quantified the time required for dorsal closure completion in these embryos, and we found a significant delay compared to controls (Figure 5I-J). Albeit unexpected, these results suggested an interdependence between epidermal and tracheal development. We decided to test an inverse scenario, where we affected dorsal closure and observed an effect on tracheal development. To do this, we used mutant embryos where epidermal dorsal closure was affected. Mutations in genes that code for components of the JNK signalling pathway either lead to a complete failure or to a delay in dorsal closure. We used *kay^1^*(which codes for Fos) mutant embryos and focused on embryos with a delayed dorsal closure phenotype to see if this would have an effect in tracheal morphogenesis. We employed *btl*>CD4::mIFP to visualize the tracheal system, together with E-Cad fused to GFP to follow the progression of epidermal dorsal closure.

Like in control embryos, where we observed that dorsal closure progressed in parallel to trunk displacement (Figure 6A-C, H-I, Video S9, left), in *kay^1^*mutant embryos with a delayed closure phenotype, tracheal trunk displacement still occurred at the same rate as epidermal dorsal closure (Figure 6D-I, Video S9, right). However, in *kay^1^* mutant embryos, tracheal trunks developed malformations that were not observed in controls, which consisted in tortuosities that changed in shape over time and mimicked movements also observed in the epidermis. Again, in this mutant condition, tracheal trunks were longer than in controls (Figure 6J).

**Figure 6.**
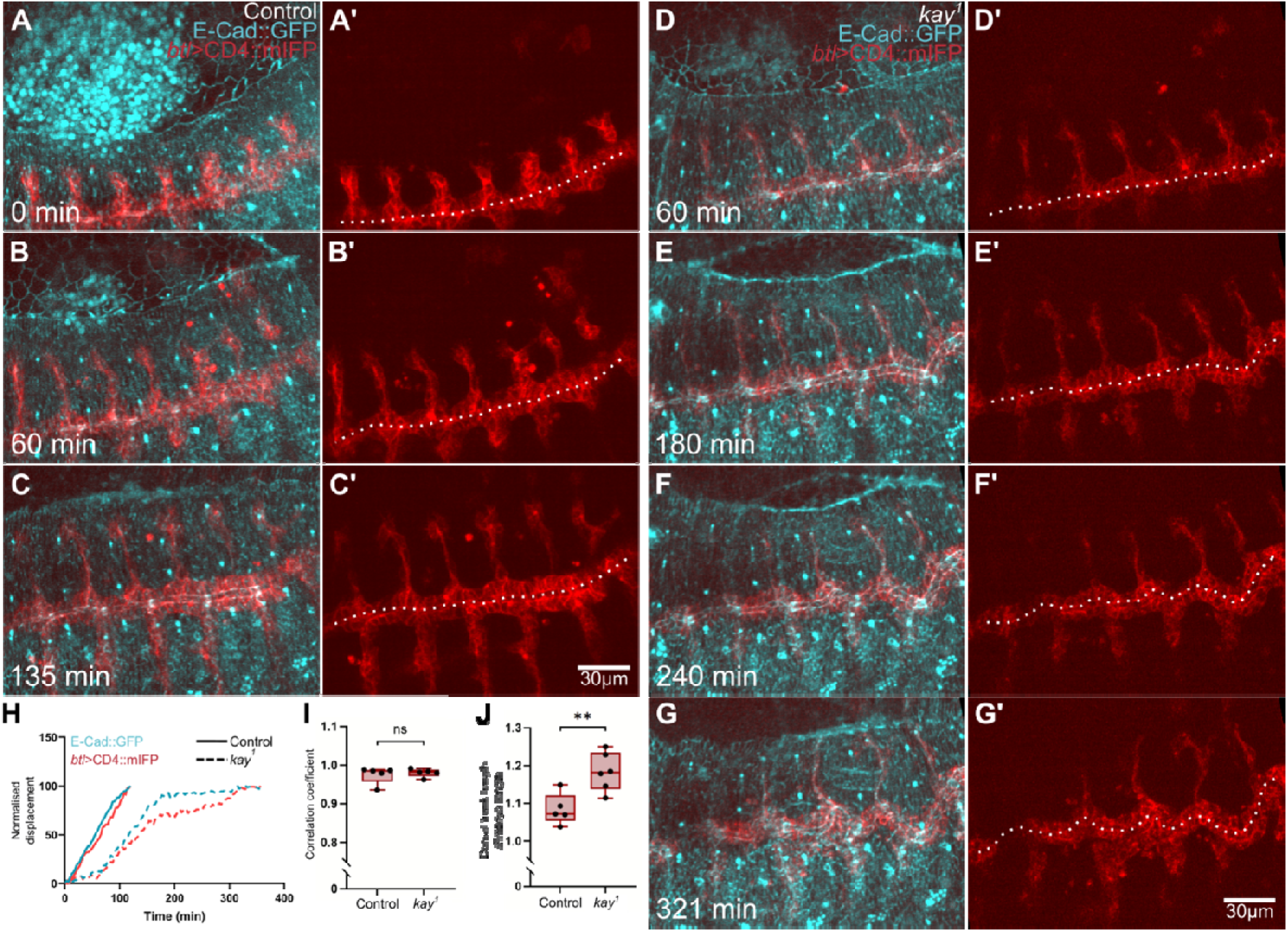
Tracheal trunk morphology in *kay^1^* mutant embryos with a delayed dorsal closure phenotype. (A-G) Maximum intensity projections and time-lapse imaging of embryos expressing E-Cad::GFP (cyan) and CD4::mIFP under *btl-gal4* (red). (A-C) Control embryo; (D-G) *kay^1^* mutant embryo. (H) Normalised displacement of the epidermis (cyan) and dorsal trunk (red). (I) Quantification of epidermis-dorsal trunk coordination, p=0.6737. Control, n=5; *kay^1^*, n=5. (J) Normalised dorsal trunk (DT) length, p=0.0058. Control, n=5; *kay^1^*, n=6. Significance was assessed using t-tests. Box plots in (I – J) represent median, IQR and max and min values.

The defects we saw on *kay* mutant embryos could be explained by a requirement for Fos in the tracheal system, however, we did HCR-FISH against *kay* in control embryos and as reported with earlier methods (Riesgo-Escovar and Hafen, 1997; Souid and Yanicostas, 2003), we detected a strong signal at the epidermal leading edge and only a faint generalized signal in the rest of the embryo (Figure S5D-D’’). To corroborate that the defects we see on the tracheal system are due to a delayed closure, we over-expressed a dominant negative form of Moesin in stripes of epidermis using *en-gal4*. In these embryos we stained for Gasp, a tracheal luminal marker, and again we found the wavy trunk phenotype (Figure S5A-C). These experiments show that the trachea-epidermis association is present even in conditions of delayed dorsal closure. Also, they support that proper trunk development depends on timely epidermal dorsal closure, and that delaying this process also delays trunk displacement.

**Figure S5.**
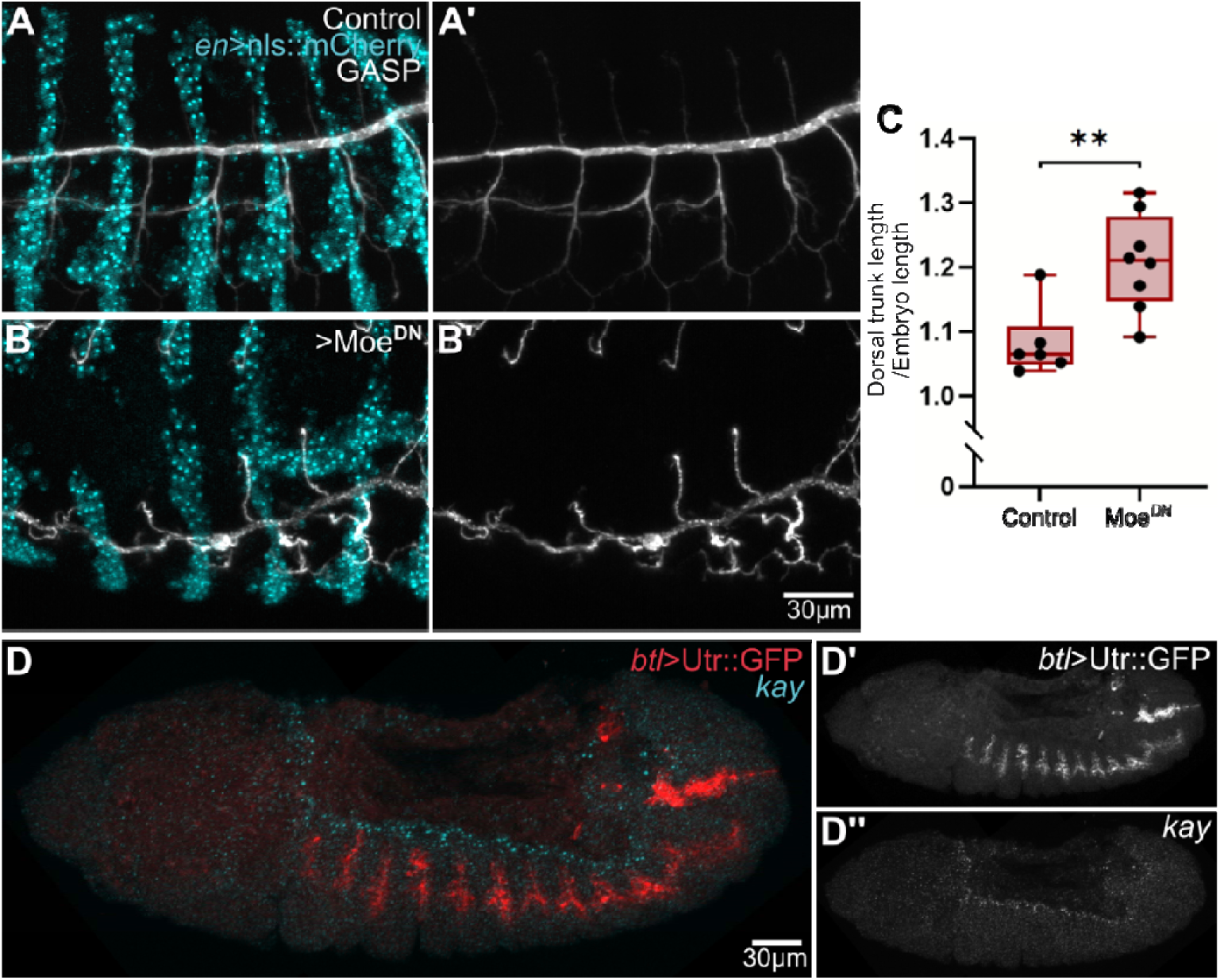
Role of the dorsal closure progression in tracheal trunk morphology. (A – B) Maximun intensity projections of embryos expressing nls::mCherry under *en-gal4* and inmunostained for GASP. (A) Control (B) Embryo with additional expression of *UAS-*Moe^DN^. (C) Quantification of normalised dorsal trunk length. Control vs Moe^DN^, p=0.0047. Significance was assessed using t-test. Box plot represents mean, IQR and min and max value. (D – D’’) HCR-FISH against *kay* (cyan) in embryos with expression of Utr::GFP under *btl-gal4* (red).

### In the absence of epidermis attachment, removing dorsal branches rescues trunk morphology

In all our manipulations, we found that trunks acquire a wavy phenotype regardless of whether we affect trachea-epidermis interactions or delay epidermal movements. In these two approaches, we think the wavy trunk phenotype may arise from different mechanisms. Upon completion of dorsal closure and as head involution takes place, the epidermis and the tracheal system elongate towards the anterior part of the embryo, so we think that in the experiments where we delay dorsal closure, the trunks grow in the antero-posterior axis while the epidermis is still moving dorsally. Since in these embryos the trunks are still attached to the epidermis, the wavy phenotype could emerge because of the mismatch in the direction of growth between the epidermis and the trunks. In fact, in *kay* mutant embryos we see that after completion of dorsal closure, the wavy trunk phenotype slightly resolves after the epidermis moves anteriorly (Video S9).

In embryos where we over-express Mmps or knock down β-Integrin, we reasoned that in the absence of or weakened contact points with the epidermis, dorsal branches could exert forces on the trunks and pull them dorsally, resulting in the wavy phenotype. We tested this by over-expressing Mmp2 and Dad simultaneously in the tracheal system. In embryos were we only over-expressed Mmp2, we found a wavy trunk phenotype like what we see when we over-expressed Mmp2 in tendon cells (Figure 7C, F), and expression of Dad rescued this phenotype (Figure 7D, F). This supports that dorsal branches can pull on the trunks as they migrate dorsally and that weakening adhesion to the epidermis leads to an uneven repositioning of the trunks.

**Figure 7.**
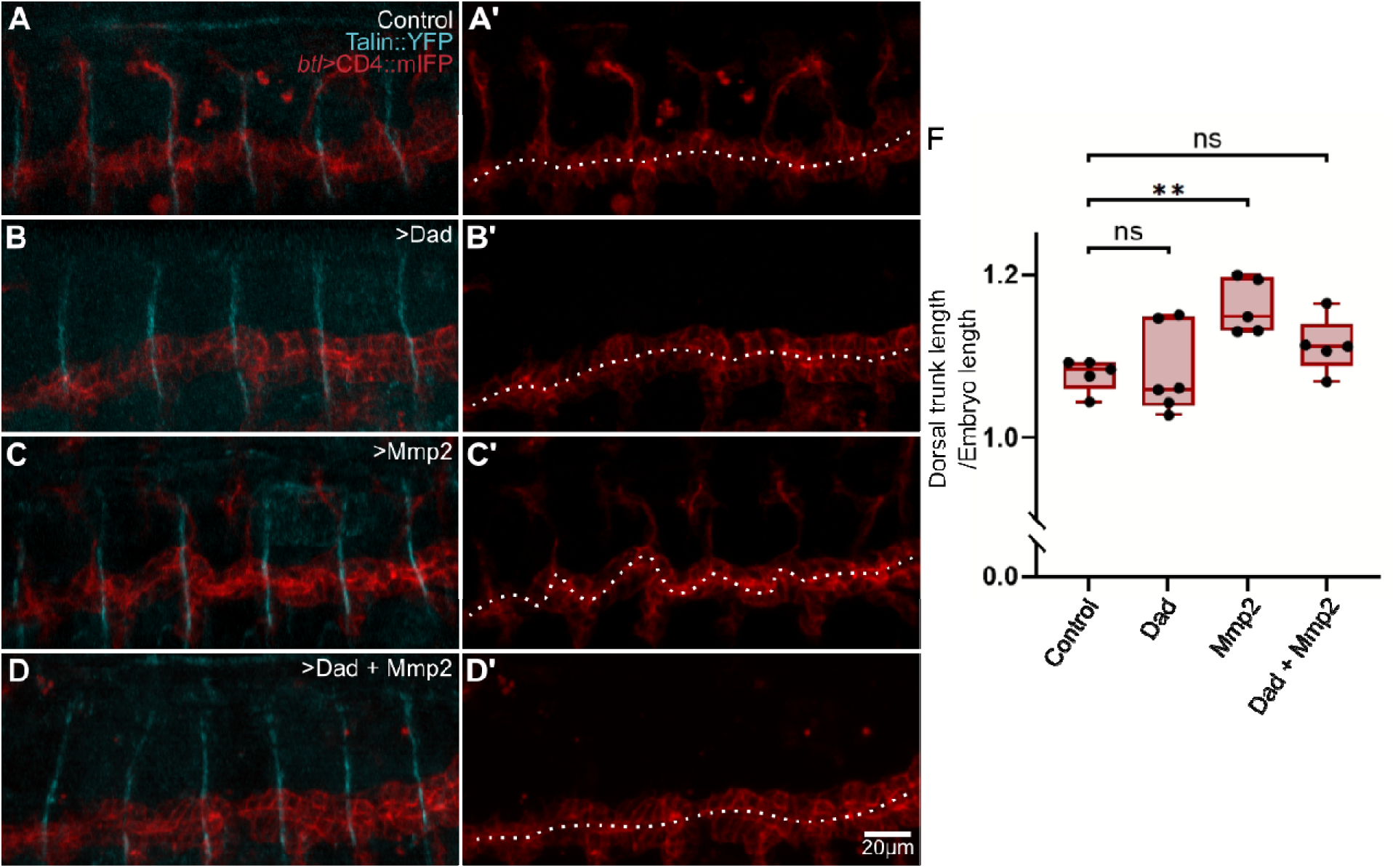
Role of the dorsal branches in the tracheal trunk malformations. (A – D) Maximum intensity projections of embryos expressing Talin::YPet (cyan) and CD4::IFP under *btl>gal4* (red). (A) Control. (B) Embryo additionally expressing UAS-Dad. (C) Embryo additionally expressing UAS-Mmp2. (D) Embryo additionally expressing UAS-Dad and UAS-Mmp2. (F) Normalised dorsal trunk length. Control vs Dad, p=0.9975; Control vs Mmp2, p=0.0088; Control vs Dad+Mmp2, p=0.3552. Control, n=5; Dad, n=6; Mmp2, n=5; Dad + Mmp2, n=5. Box plot represents median, IQR and min and max value. Significance was determined using ANOVA and Dunnett correction for multiple comparisons.

Altogether, we conclude that adhesion of the dorsal trunks to the tendon cells of the epidermis through the ECM is important for trunk repositioning. In addition, this interaction plays a stabilising role in maintaining the straightness of dorsal trunks during their movement to the dorsal side of the embryo. These interactions are important for tracheal morphogenesis, but they also have an effect in epidermal dorsal closure.

## Discussion

Here, we demonstrated that epidermal closure and tracheal trunk displacement are coordinated events, with evidence suggesting that ECM integrity plays a key role in their morphogenesis. In this model, the ECM would act at the same time as a milieu where extracellular signals can diffuse and act in different tissues, and as a scaffold that transmits forces, linking tissues as they undergo complex rearrangements. In the case of trachea-epidermis interactions, a number of classical studies have shown that signals produced by the epidermis act in the epidermis itself but also regulate tracheal patterning [(Affolter et al., 1994; Chihara and Hayashi, 2000; Glazer and Shilo, 2001; Kato et al., 2004; Llimargas, 2000), reviewed in (Sánchez-Cisneros et al., 2025)]. Our experiments show that besides these well-characterised signals, mechanical interactions between the two tissues also fine-tune tracheal behaviour. We show that pulling forces from dorsal tip cells are dispensable for trunk displacement, but they are still able to pull on the trunks when these are not bound to the epidermis. Instead, trunk displacement is facilitated by adhesion to the epidermis, and this interaction also influences epidermal dorsal closure (summarized in Figure 8).

**Figure 8.**
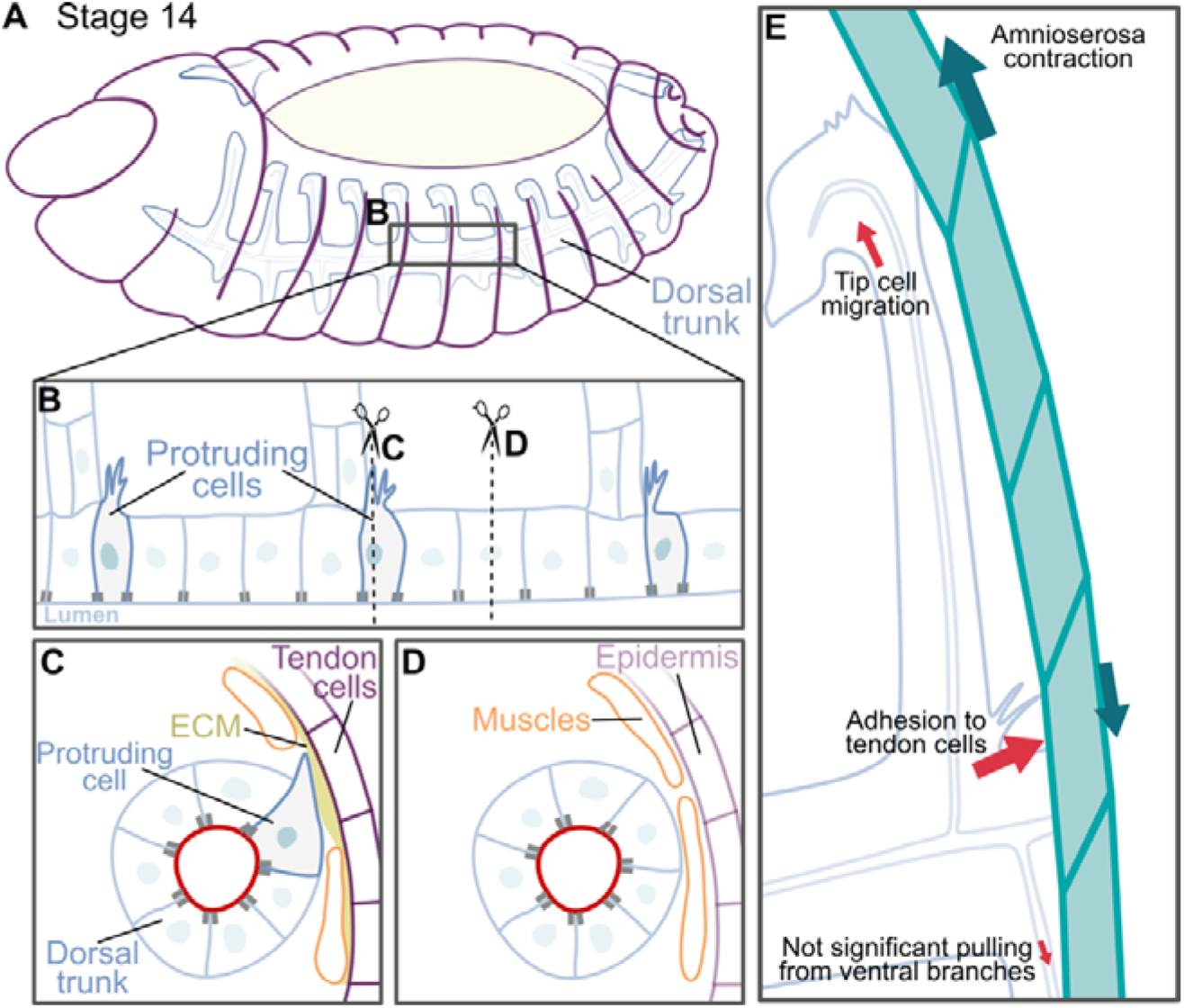
Model of trachea-epidermis interactions. (A) Overview of a stage 14 embryo at the onset of dorsal closure. The black square at the dorsal trunk is magnified in (B). (B) Protruding cells distribution at the dorsal trunk. The dotted lines are presented as cross-sections in (C) and (D). (C) Just below tendon cells, protruding cells extend filopodia towards the basement membrane of tendon cells. (D) Regions of the trunks not near to tendon cells are separated from the epidermis by muscles and other mesenchymal cells. (E) Summary of forces acting during dorsal trunk displacement. The interactions between the protruding and tendon cells allow trunk repositioning at the same time as they counteract the pulling forces produced by migrating tip cells. In addition, during this process, the dorsal trunk acts as a counterweight for the epidermis, balancing epidermal dorsal closure progression.

### Emergence of a ‘wavy trunk’ phenotype upon disruption of trachea-epidermis interactions

Our results demonstrate that perturbing trachea-epidermis interactions by different mechanisms (weakening the ECM through Mmp over-expression, targeting tracheal β-integrin to degradation, or delaying epidermal dorsal closure), consistently result in similar phenotypes: trunks are still displaced dorsally but they acquire a wavy appearance. This suggests that tracheal trunk displacement is in fact dependent on epidermal development. Interactions between trunks and the epidermis should allow trunk repositioning but also prevent deformation by the pulling forces that result from dorsal branch elongation. A previous work that focused on dorsal branch cell development used laser micro-dissection to show that tip cell migration is responsible for branch elongation and intercalation, which makes likely that force generated by tip cells can act on the trunks, at the base of dorsal branches (Caussinus et al., 2008).

The wavy trunk phenotype has been already reported as a result of loss of laminins (Klußmann-Fricke et al., 2022). Despite the phenotype that manifests in laminin (*LanB1*) mutant embryos is stronger than the ones that we report here, we interpret it at least partly because of the loss of the interaction with the epidermis. The stronger phenotype described in laminin mutant embryos could be explained by an additive effect of cell-autonomous defects in cell polarity, as previously suggested (Klußmann-Fricke et al., 2022), by an incomplete silencing of integrin-mediated adhesion in our experiments, and by the presence of other adhesion complexes that rely on laminin besides the integrins complex.

### Interactions between protruding cells and tendon cells

According to our model, trachea-epidermis interactions occur at specific contact points distributed along the antero-posterior axis of the embryo, where tracheal trunks intersect with tendon cells of the epidermis. The protruding cells we describe here are frequently found directly in front of tracheal dorsal branches, which we believe has prevented their characterisation as they are difficult to visualise with general tracheal reporters.

Tendon cells are well-known force transmitters as they interact with the exoskeleton and muscle cells through apical and basal ECM layers, respectively, enabling locomotion as muscles contract (Chu and Hayashi, 2021; Martin-Bermudo, 2000; Weitkunat et al., 2014). Our analyses suggest that these points also mediate the adhesion of tracheal trunk cells, the protruding cells, to transmit forces relevant for tracheal remodelling. As mentioned earlier, in face of pulling forces exerted by dorsal tip cell migration, adhesion to the tendons may counteract trunk deformation. Through this interaction, stalk cells can intercalate as trunks remain in place. (Caussinus et al., 2008; Ochoa-Espinosa et al., 2017). We do not see defects in cell intercalation in the conditions we tested, but these could be compensated by the looping of the trunks upon pulling interactions from tracheal branches.

Our results do not address if protruding cells possess different gene-expression profiles compared to their neighbours or if they acquire a distinctive morphology due to their proximity to the specialised ECM found near tendon cells. Previous works have shown that tendon cells secrete FGF which in turn regulates tracheal development (Butí et al., 2014; Dorfman et al., 2002). However, it is unlikely that tendon cell-derived FGF is responsible for the formation of these protrusions, as tracheal trunk cells do not activate landmarks of FGF signalling like SRF expression or ERK phosphorylation (Gervais and Casanova, 2011). Previous studies have induced tendon cell differentiation by over-expression of Stripe in other epidermal cells (Dorfman et al., 2002). Because of the effect on FGF expression, it is difficult to assess the effect on trunk cell behaviour from these experiments.

### Trachea-epidermis interactions and their role in epidermal dorsal closure

Besides the effects on the trunks, we found that targeting β-integrin in the tracheal system also resulted in a delay in epidermal dorsal closure. Previously, it was shown that dorsal tip cell migration is independent from epidermis displacement (Caussinus et al., 2008). Our results do not contradict these experiments; we see adhesion only at the level of tracheal trunks, and from our experiments there are no effects on tip cell morphology or migration. It has been shown that the forces produced by the amnioserosa are the main components that allow epidermal dorsal closure; even when the epidermis forms a supracellular actomyosin ring at dorsal-most row of epidermal cells, this only controls the seamless sealing of the epidermal sheets at the dorsal midline (Pasakarnis et al., 2016). It was suggested recently that the epidermis itself acts as a counter-force in response to the pulling forces of the amnioserosa (Lv et al., 2022). Our results support this view and suggest that adhesion of the trunks to the epidermis may contribute to balance the interplay of forces that participate in epidermal dorsal closure. From removing a force that opposes epidermal stretching, we did not anticipate a delayed dorsal closure. However, the delay could be explained by a disorganization in the movements of the epidermis. In accordance, mutants for genes that negatively regulate dorsal closure progression also affect proper sealing of the epidermis (Jacinto et al., 2002; Martín-Blanco et al., 1998).

### The ECM as a coordinator of morphogenesis

Mechanical interactions between adjacent tissues and how they shape morphogenesis have been described in a wide range of processes. These include from the above-mentioned epidermis-amnioserosa interactions in *Drosophila* development (Fernández et al., 2007; Frank and Rushlow, 1996; Hayes and Solon, 2017; Kiehart et al., 2017) to epidermis-blood vessels in the rhinarium of dogs and other species (Dagenais et al., 2024). In many of these processes, the ECM, its mechanical properties and its remodelling, play important roles in force transmission and/or coordinating tissue behaviour [(Goodwin et al., 2016; Goodwin et al., 2017; Inoue and Hayashi, 2007; Münster et al., 2019), reviewed in (Barrera-Velázquez and Ríos-Barrera, 2021)]. We suggest that ECM-mediated tissue adhesion may have a role in general embryo development, integrating morphogenetic processes of seemingly unrelated tissues.

## Author contributions

Conceptualization, LDRB; Methodology and investigation, LESC, MBV, DK, HML, LDRB; Formal Analysis, LESC, MBV, PB, LDRB; Writing-first draft, LESC & LDRB; Writing—Review and Editing, LESC, MBV, DK, PB, HML, LDRB; Visualization, LESC & LDRB; Supervision, LDRB; Funding Acquisition, LDRB.

## Acknowledgements

We thank the Ríos-Barrera lab and Sergio Romero-Romero for critical discussions and feedback on the manuscript. We also thank the technical support of Alejandro Marmolejo-Valencia (Merchant-Larios lab), Berenice Otero-Díaz (Ríos-Barrera lab) and Miguel Tapia-Rodríguez (Unidad de Microscopía, Instituto de Investigaciones Biomédicas, UNAM, Mexico). We acknowledge Euro-BioImaging ERIC (https://ror.org/05d78xc36) for providing access to imaging technologies and services via the EMBL Node in Heidelberg, Germany (https://ror.org/03mstc592), and we thank Stefan Terjung (Advanced Light Microscopy Facility, EMBL) for his support. LESC is a student from the program Doctorado en Ciencias Bioquímicas and is funded by a CONAHCyT/SECIHTI fellowship CVU #1099237. This work was supported by Universidad Nacional Autónoma de México, Programa de Apoyo a Proyectos de Investigación e Innovación Tecnológica (UNAM-PAPIIT) grant #IA202923 and an Early Career Return grant from the International Centre for Genetic Engineering and Biotechnology, Italy, #CRP/MEX21-04_EC.

## Notes

### Competing Interest Statement

The authors have declared no competing interest.

### Summary of Updates

We incorporated kind suggestions from three anonymous reviewers, which we received through Review Commons. The modifications have been listed in the revision plan posted along with our manuscript and briefly include: - Experiments that show the effect of the over-expression of UAS-Dad in trunk displacement - Experiments where epidermal dorsal closure was affected by the over-expression of a dominant negative form of Moesin under engrailed-gal4 - Updates on a summary figure where we incorporated a model of our work - Other changes in the text and incorporation of references to original research papers instead of reviews

